# GLYCO-2: a Tool to Quantify Glycan Shielding of Glycosylated Proteins with Improved Data Processing and Computation Speed

**DOI:** 10.1101/2025.02.26.640297

**Authors:** Mateo Reveiz, Myungjin Lee, Peter D. Kwong, Tongqing Zhou, Reda Rawi

## Abstract

**Motivation:** Glycans bound to glycoproteins mediate immune response, including antibody recognition and immune evasion. Previously, we developed an *in silico* tool GLYCO (GLYcan COverage) to quantify the glycan shielding of a protein surface, applying it to various studies. However, GLYCO lacks sufficient computational efficiency when analyzing larger datasets.

**Results:** Here we introduce GLYCO-2 which improves the computational speed by ∼4- fold compared to GLYCO by adopting a new analytical cylinder method with *k*-d trees. GLYCO-2 can calculate glycan shielding from a single coordinate file or from multiple frames derived from molecular dynamics simulations accounting for the inherent flexibility of oligosaccharides. We applied GLYCO-2 to quantify glycan shielding of influenza hemagglutinin (HA) proteins across diverse subtypes that infect humans, revealing an increasing trend in glycan shielding over time within each subtype, likely contributing to immune evasion. Overall, the enhanced computational efficiency of GLYCO-2 allows for faster and easier quantification of glycans, which contributes to the understand of glycan shielding effects in fields such as immunology and vaccine design.

**Availability and implementation:** GLYCO-2 is freely available at https://github.com/meteosR/GLYCO-2/

**Supplementary information:** Supplementary data are available at Bioinformatics online.

## 1 Introduction

Glycans, also referred to as polysaccharides, serve as pivotal regulators of various biological processes. These complex moieties play a fundamental role in structural stabilization and contribute to crucial functions, including the recognition of antibodies by antigens (Kang, et al., 2020; Lee, et al., 2021; More, et al., 2016; Wei, et al., 2020). Glycans are complex assemblies of sugar moieties. The complexity of glycans comes from their intricate molecular structures, characterized by diverse arrangements of monosaccharides and branching patterns. They show a wide range of variations in size, composition, and linkage, contributing to their remarkable heterogeneity, which can be elucidated through site-specific glycan analysis as demonstrated in recent studies of viral glycoproteins (Struwe, et al., 2018; Watanabe, et al., 2020).

The complexity and inherent flexibility of glycans not only necessitates advanced method for accurate quantification but also highlights the significance of glycan research in the biological systems. In this context, understanding the structural and functional aspects of glycans is significant for advancing our knowledge to develop innovative therapeutic strategies.

Previously, we developed an *in silico* tool, GLYCO (GLYCO-1) (Lee, et al., 2021) to quantify glycan coverage of glycoproteins. It has been utilized in many works such as correlation with glycan coverage and antigen framework antibody distance (Lee, et al., 2021), glycan density evaluation of N-terminus domain supersite of SARS-CoV-2 (Cerutti, et al., 2021), quantification of glycan density on *O*-linked and *N*-linked glycosites on V3 loop of SIV (Gorman, et al., 2022). However, GLYCO-1 requires extensive computation time for glycan quantification, particularly when analyzing multiple structural coordinate files, such as those generated from molecular dynamics simulations covering hundreds of nanoseconds to microseconds (Casalino, et al., 2021).

Here, we introduce GLYCO-2, an improved version of GLYCO-1, that implements a mathematically more robust algorithm with more efficient computation performance. With the accelerated version, rapid analysis of massive numbers of PDB files becomes available within a concise timeframe. As case studies, GLYCO-2 was used to analyze influenza hemagglutinin proteins and their glycan shields.

## 2 Methods

### 2.1 GLYCO-2 algorithm

GLYCO-2 improves glycan evaluation by using an analytical solution by efficiently querying for protein coordinates with *k*-d trees. Glycan coverage was defined as the number of glycan atoms within a cutoff centering from a surface protein residue except the ones that are hindered by protein region. The evaluation of hindered glycan atoms was examined by having a vector between the surface protein residue to each glycan atom. Previously, GLYCO-1 used a series of cubic boxes (numerical line parameterization) to detect blocking protein atoms along the vector (Figure 1A left). However, in GLYCO-2 we replaced the chain of cubic boxes with a single cylinder (analytical cylinder method) to evaluate if glycan atoms shield a corresponding surface protein atom (Figure 1A right).

**Figure 1.**
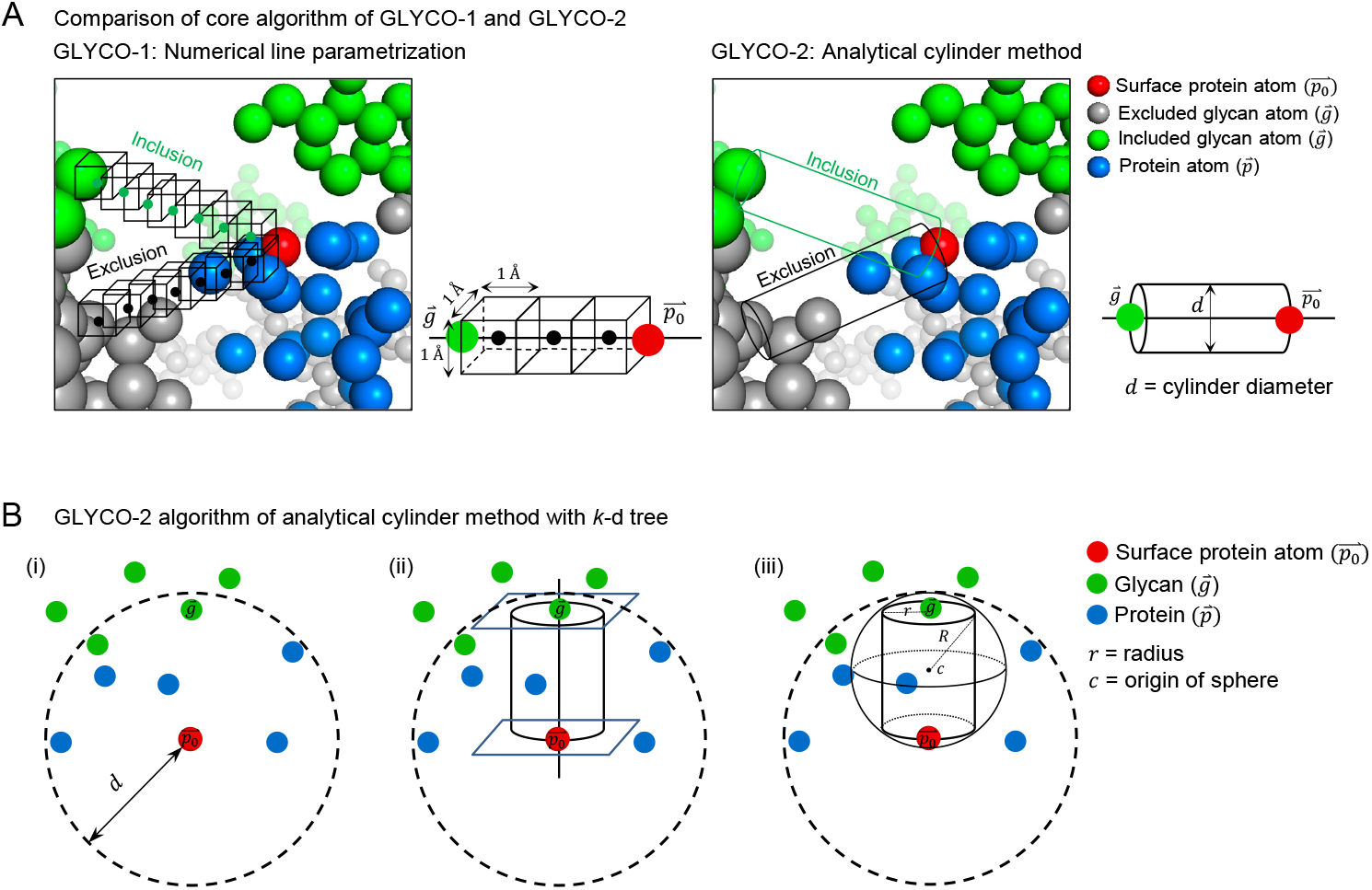
GLYCO-2 algorithm refines the shielding definition. (A) General comparison between methods GLYCO-1 and GLYCO-2. (B) The core algorithm of GLYCO-2 selects shielding glycans for surface protein atoms using an analytical cylinder method with k-d trees.

More specifically, let 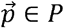 be a protein atom and 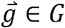 be a glycan atom. Let *G*_*p*_ ⊆ *G* be the set of glycan atoms within cutoff distance, *d* of 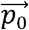 (Figure 1B-i):

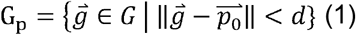

We say that a glycan atom 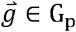 shields 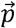 if there is no other protein atom in between them. In other words, if there is no protein atom *p* inside the cylinder defined by *g* and *p*_*0*_ at the center of the bases and a radius *r* (Figure 1B-ii).

Shielding of 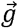 to 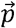 denoted *S* (*g,p*) can be calculated as follows:

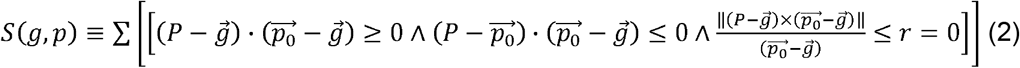

Glycan coverage for a given amino acid x is then the sum over all the glycan atoms in *G*_*p*_ across all the heavy atoms (excluding hydrogen atoms) conforming the residue A.

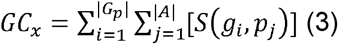

Second, GLYCO-2 adopts efficient algorithm to retrieve protein atom coordinates, which improve computational speed. The *P* term in equation (2) can be further optimized by noting that only a small subset of atoms *P*_*S*_ ⊆ *P* inside the cylinders’ inscribing sphere of origin *c* and radius R could be contained within the cylinder (Figure 1B-iii). The subset *P*_*S*_ can be obtained by querying a precomputed *k*-d tree containing all protein atom coordinates, as follows:

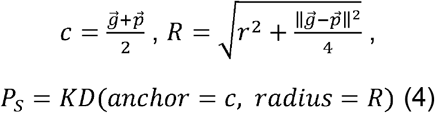

In practice, we find that *P*_*S*_ ≪ *P* if the threshold distance d remains small (default value is 23 Å) and most glycans are found on the protein surface.

### 2.2 Dataset: glycosites of influenza virus hemagglutinin

Influenza A subtype H3N2 and H1N1 sequences from 1968 (1918 for H1N1) to 2024 were collected from NCBI Virus, Bethesda (MD): National Library of Medicine (US), National Center for Biotechnology Information; [cited 2024 08 28]. Available from: https://www.ncbi.nlm.nih.gov/labs/virus/. Only human full-length available sequences were chosen as input dataset. Glycan sequons per sequence were collected by netNglyc-1.0 (Gupta and Brunak, 2002). Hemagglutinin sequences per year were clustered via CD-HIT (Fu, et al., 2012; Li and Godzik, 2006) with sequence identity threshold 97%.

### 2.3 Structure modeling

The representative sequence per cluster was modeled using YASARA (Krieger and Vriend, 2014). Homology modeling was performed for these target sequences, utilizing five templates automatically selected by YASARA. Mannose-5 glycans were modeled on all glycosites using Glycosylator (Lemmin and Soto, 2019).

### 2.4 GLYCO-2 usage

The user can enter one or multiple PDB coordinate files with glycosylated molecules built from glycosylation programs such as CHARMM-GUI Glycan Reader & Modeler (Jo et al., 2011; Park et al., 2017; Park et al., 2019) or Glycosylator (Lemmin and Soto, 2019). Glycan coverage for user defined polysaccharides can be computed for the entire surface of protein or a given subset of residue-only by selecting module type. Also, users are free to enter parameters such as solvent accessible surface area, glycan distance cutoff, and cylinder radius as described in GLYCO-1 (Lee et al., 2021). The output files contain the glycan coverage with corresponding protein/glycan details and the glycan coverage overlayed PDB file. Glycan coverage overlayed file stores the quantified values in the B-factor column that can be displayed by molecular visualization programs highlighting dense and sparse glycosylation regions.

## 3 Results

### 3.1 GLYCO-2 provides rigorous solution and improves computational speed compared to GLYCO-1

The use of an analytical definition of a cylinder coupled with *k*-d trees greatly boosts the computational speed of glycan quantification. In practice, 20 distinct datasets containing different protein geometries, glycan types (*N*-linked and *O*-linked) and glycosylation patterns from SARS-CoV-2 spike, HIV-1 Env, Ebola, Zika, SIV and influenza HA were used to benchmark the performance of both GLYCO versions (Supplementary Table 1). Glycan coverage values for the two versions are almost identical with marginal differences (R^2^ = 0.97). GLYCO-1 slightly overestimates glycan coverage by including more glycan atoms in the calculation when compared to a cylinder radius of 1.4 Å for GLYCO-2 (Figure 2A). GLYCO-2 showed an average ∼4-fold computation speed improvement for both single and multiprocessing settings (Figure 2B). The new algorithm also showed higher degrees of parallel efficiency. An efficiency of 0.74 was achieved with 8 CPUs when using Intel(R) Xeon(R) Gold 6154 CPU @ 3.00GHz models on Ubuntu 18.04 (Figure 2C).

**Figure 2.**
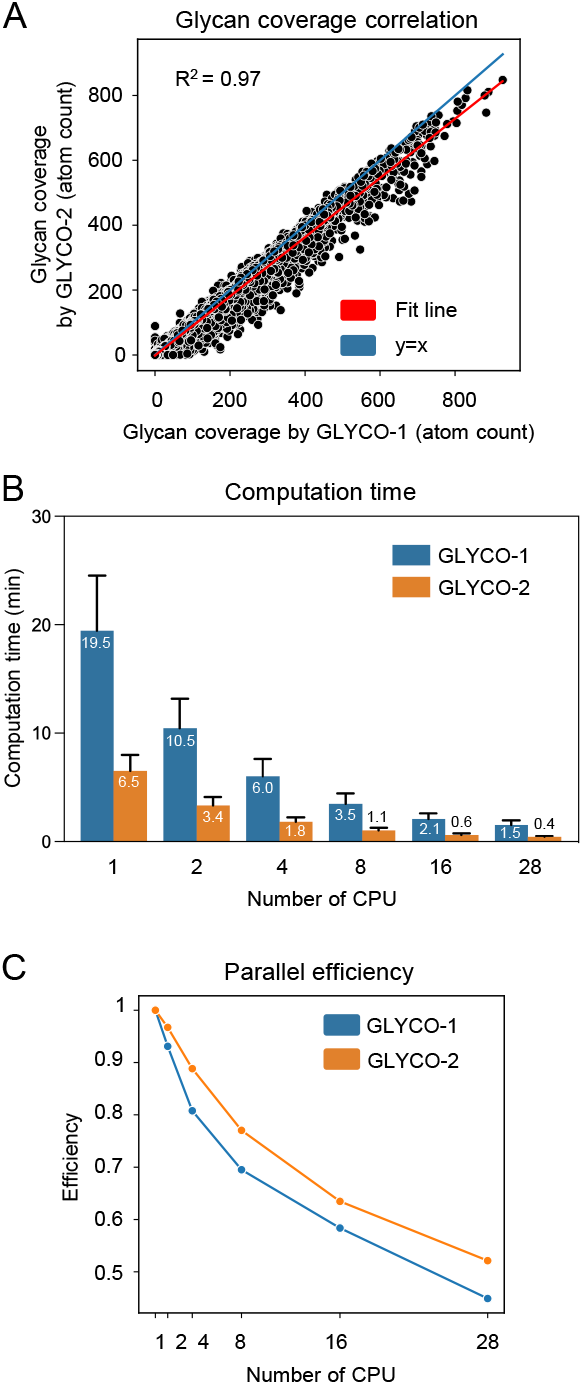
GLYCO-2 computationally outperforms its previous version, GLYCO-1. (A) Correlation of glycan coverage between GLYCO-1 and GLYCO-2 across 20 aggregated datasets, encompassing diverse glycoproteins under multiple running settings. (B) Comparison of average computational times between GLYCO-1 and GLYCO-2, with the average labeled on the plot and standard deviation indicated by black bars. (C) Comparison of parallel efficiency between GLYCO-1 and GLYCO-2.

### 3.2 Benchmark study

#### 3.2.1 The effect of parameter variation

We further examined the impact of altering parameters – cylinder radius, glycan distance cutoff, and solvent accessible surface area.

The default cylinder radius 1.4 Å was defined as twice the atomic radius of a carbon atom (0.7 Å) to completely exclude unintended protein shielding (Figure 3A, left). We evaluated the effect of changes in cylinder radius on the estimation of glycan coverage. Given that larger cylinder radius excludes more glycan atoms, glycan coverage decreases as the cylinder radius increases from 0.2 to 2.6 (Figure 3A, middle and right).

**Figure 3.**
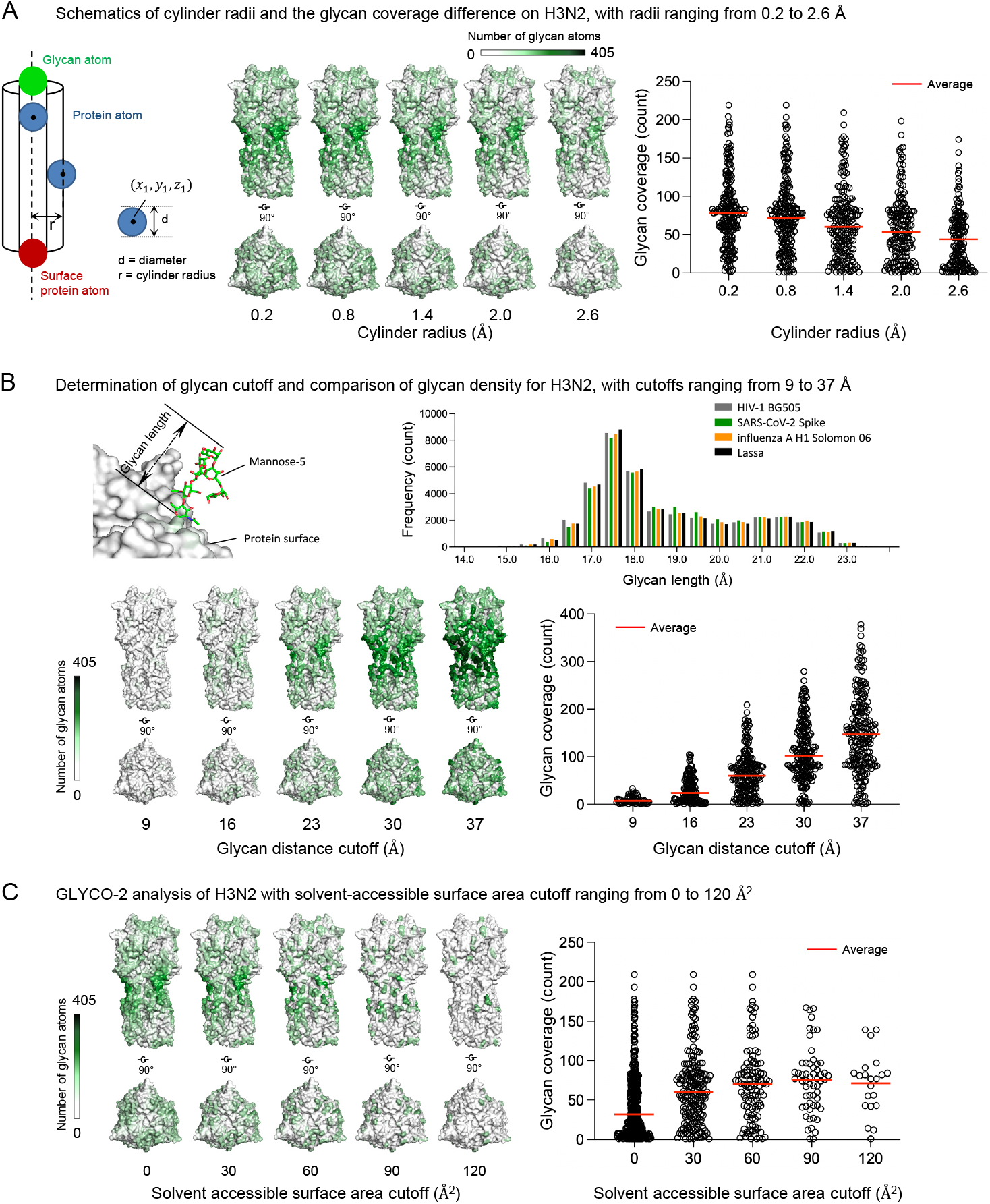
Change in glycan coverage of Influenza A H3N2 with varying parameters. Glycan coverage changes as cylinder radius (A), glycan distance cutoff (B), and solvent-accessible surface area (C) varies. A glycan distance cutoff of 23 Å, surface area cutoff of 30 Å^2^, and cylinder radius of 1.4 Å were used when these parameters were not being examined.

In previous work (Cerutti, et al., 2021; Lee, et al., 2021) we used a glycan distance cutoff as defined by the longest length of glycans (23 Å) throughout MD trajectories for mannose-5 glycosylated HIV-1, SARS-CoV-2, influenza A and Lassa virus (Figure 3B, top two panels). The idea was to consider the maximum available glycan shield length that a single mannose-5 can potentially assume. However, users can alter the glycan distance cutoff depending on their systems. A low cutoff can depict the local glycan covering effect, whereas a long cutoff, such as longest length of glycan, provides maximum glycan coverage as possible. Clearly, as the cutoff increases, more glycans are included, resulting in dense coverage on protein surface (Figure 3B, bottom two panels).

Glycan coverage calculations require the definition of surface atoms. A reference cutoff of 30 Å^2^ is commonly used to select surface atoms, but users can adjust threshold based on their analysis needs. Lowering the cutoff includes more buried protein atoms in the calculation, allowing for an assessment of their associated glycan coverage (Figure 3C, left). However, because buried protein atoms are farther from glycans, they experience less shielding. Conversely, increasing the cutoff selects only exposed protein atoms, excluding buried atoms but resulting in higher glycan shielding (Figure 3C, right).

#### 3.2.2 GLYCO-2 proves glycan accumulation on globular head of influenza hemagglutinin H3N2 over years

We analyzed glycan coverage of influenza HA protein of subtype H3N2 circulating from 1968 to 2024 using GLYCO-2 as a case study. Glycan analysis confirms the previously observed glycan accumulation since the pandemic year of 1968 on the globular head of HA, proximal to the sialic acid binding site (Figure 4A and 4B). Given sialic acid being a primary receptor of HA, glycan coverage analysis clearly proves that HA evolves by accumulating more glycans on the head to evade immune response. This becomes clearer by observing glycan coverage for head and stem, separately (Figure 4C). Glycan coverage on head shows an upward trend (black) along with the increasing number of glycosites (red), whereas glycan coverage on the HA stem remains nearly constant, with no additional glycosites.

**Figure 4.**
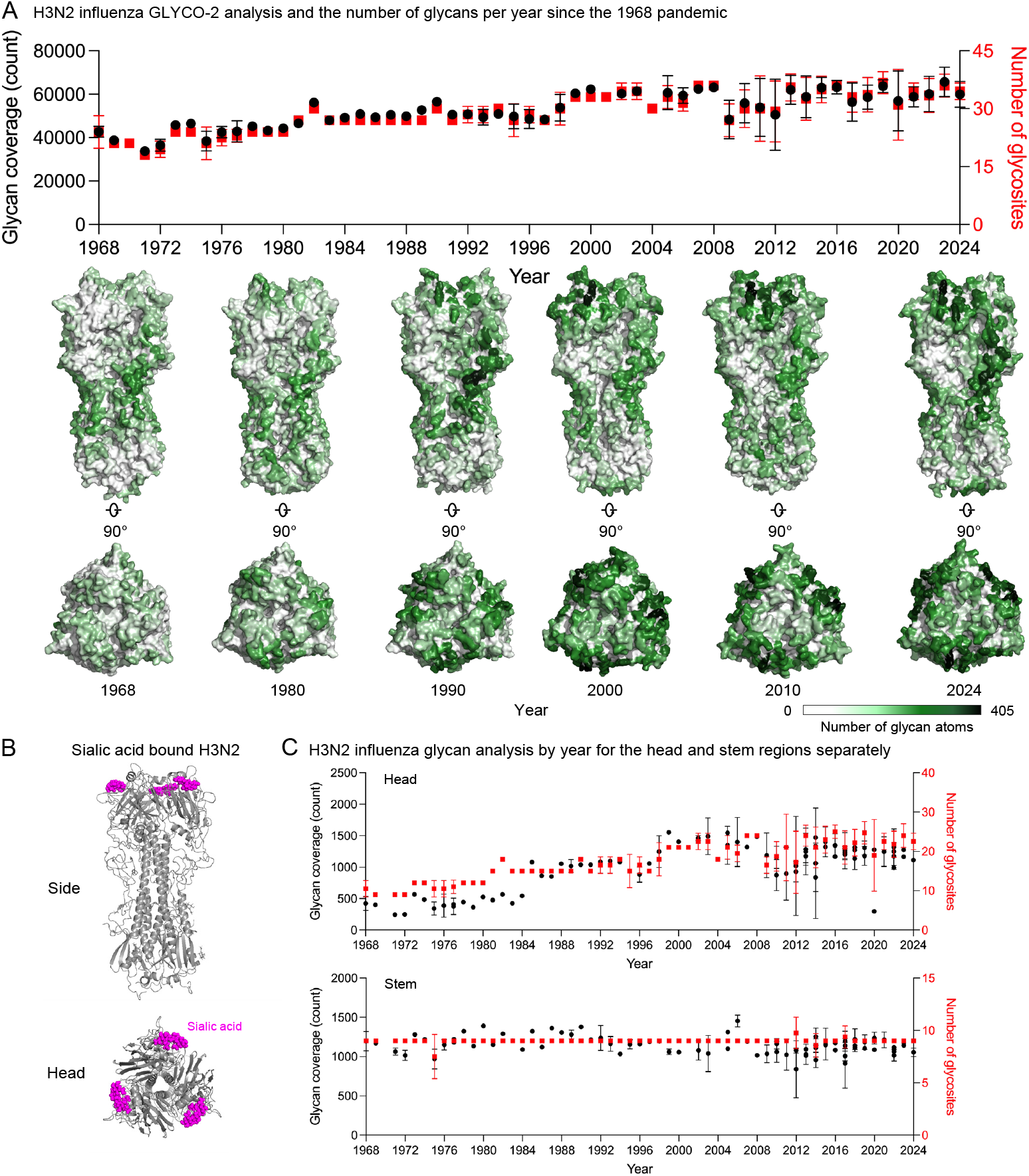
Benchmark study of GLYCO-2.0 to analyze glycan density of mannose-5 glycosylated influenza hemagglutinin H3N2. (A) Glycan coverage analysis by GLYCO-2.0 (black) and number of glycan sequons (red) for influenza hemagglutinin H3N2 from 1968 to 2024. The corresponding glycan coverage overlaid structures are shown below the plot. A glycan distance cutoff of 23 Å, a surface area cutoff of 30 Å^2^, and a cylinder radius of 1.4 Å were used to run GLYCO-2.0. (B) Sialic acid-bound H3N2 (PDB: 1HGG) viewed from the side and top. Sialic acids are represented as spheres and colored magenta. (C) Head and stem-separated glycan coverage by GLYCO-2.0 (black) and the number of glycan sequons (red) for influenza hemagglutinin H3N2.

A similar analysis of H1N1 influenza reveals distinct evolutionary patterns (Supplementary Figure 1). Glycan coverage trends from 1918 to 1958 show an initial rise, followed by a sharp decline with the emergence of H2N2, which dominated from 1957 to 1968. H1N1 resurfaced in 1975, marking its reappearance in the viral population. Notably, glycan coverage on the head exhibits a distinct pattern, with accumulation observed from 1918 to 1957, followed by a continuous trend until a noticeable drop in 2009.

## Conclusion

We developed a powerful glycan quantification tool GLYCO-2. We dramatically improved the performance compared to the previous iteration, by upgrading the key glycan evaluation algorithm integrated with *k*-d tree method. Consequently, analyzing multiple frames from MD simulations are available to inspect inherent flexibility of oligosaccharides with short amount of computational time.

Beyond version 2, we plan to further update the program to enhance its biological impact by correlating glycan coverage with other immunogenic properties. With this update, the revised program will provide improved insight into glycan coverage and immunology.

## Acknowledgements

We thank J. Stuckey’s assistance for manuscript editing. We also thank members of the Structural Virology and Vaccinology Section and Structural Bioinformatics Core, Vaccine Research Center, for discussions on the manuscript. This work utilized the computational resources of the NIH HPC Biowulf cluster.

