## Supplemental Material for "GLYCO-2: a Tool to Quantify Glycan Shielding of Glycosylated Proteins with Improved Data Processing and Computation Speed"

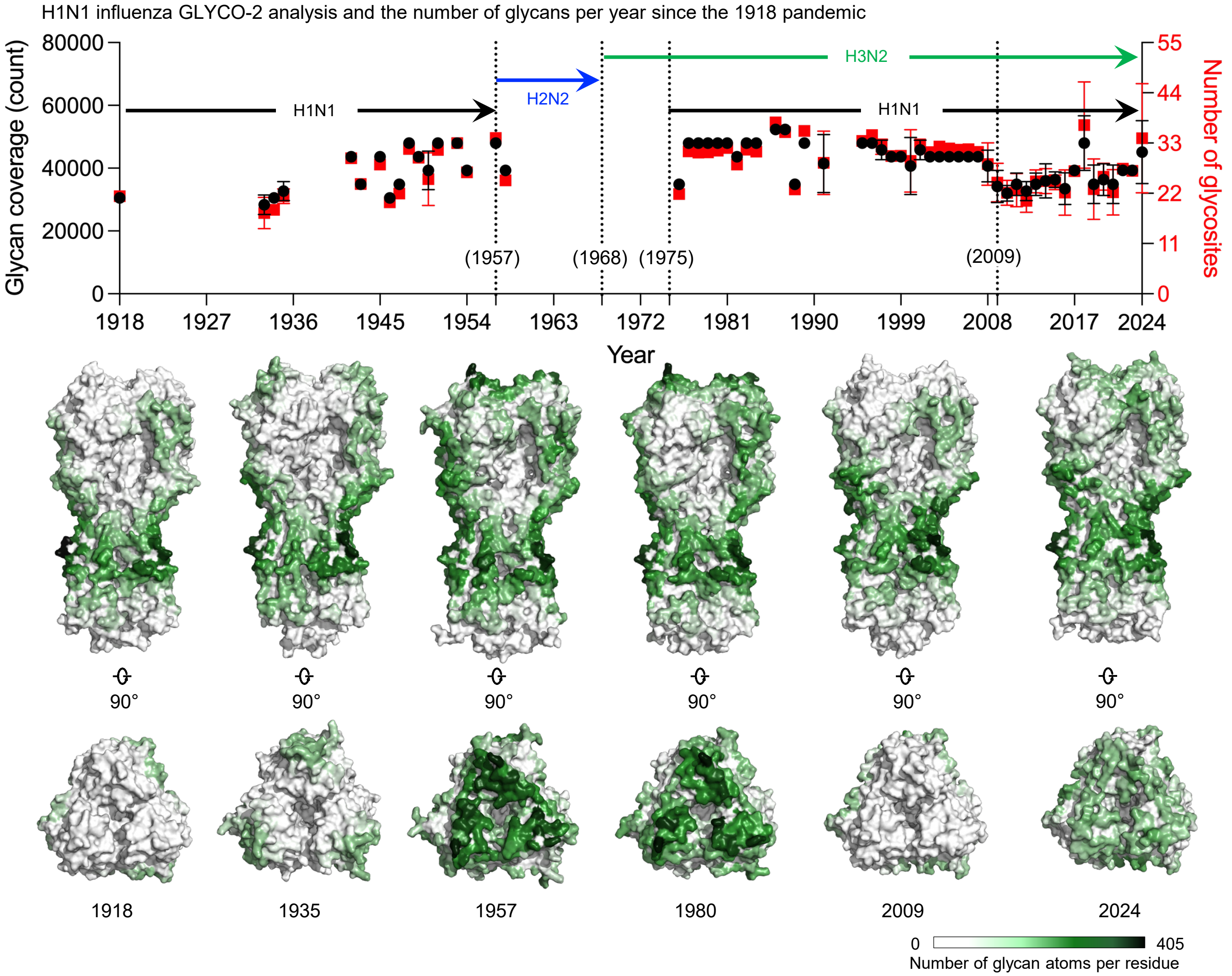


**Figure S1. Benchmark study of GLYCO-2 to analyze glycan density of Influenza A H1N1.**
Glycan coverage (black) and sequon (red) analysis for Influenza A H1N1 from 1918 to 2024. Corresponding glycan coverage overlaid structures are shown below the plot. A glycan distance cutoff of 23 Å, a surface area of 30 Å², and a cylinder radius of 1.4 Å were used for the calculations. Pandemic periods of H3N2, H2N2, and H1N1 are marked with arrows.

**Table S1. 20 test cases for comparing GLYCO-1 and GLYCO-2.**

| | Viral antigen | PDB | Glycan | Glycan distance cutoff | Surface area | Cylinder radius |
|---|---|---|---|---|---|---|
| 1 | Ebola | One of PDBs from a MD trajectory | *N*-linked mannose-5 (BGLN, BMAN, AMAN) | 23 | 30 | 1.4 |
| 2 | Ebola | 3S88 | *N*-linked mannose-5 (NAG) | 10 | 30 | 1.4 |
| 3 | HIV-1 Env trimer | One of PDBs from a MD trajectory | *N*-linked mannose-5 (BGL, BMA, AMA) | 23 | 30 | 1.4 |
| 4 | HIV-1 Env trimer | One of PDBs from a MD trajectory | *N*-linked mannose-9 (BGL, BMA, AMA) | 26 | 30 | 1.4 |
| 5 | HIV-1 Env trimer | One of PDBs from a MD trajectory | Site specific (AFU, BGL, AMA, BGA, ANE) | 26 | 30 | 1.4 |
| 6 | HIV-1 Env trimer | One of PDBs from a MD trajectory | *N*-linked mannose-5 (BGL, BMA, AMA) | 23 | 30 | 1.4 |
| 7 | HIV-1 Env trimer | One of PDBs from a MD trajectory | *N*-linked mannose-5 (BGL, BMA, AMA) | 23 | 30 | 1.4 |
| 8 | HIV-1 Env monomer | 5FYL – monomer | Incomplete glycans in PDB (NAG, BMA, AMA) | 10 | 30 | 1.4 |
| 9 | HIV-1 Env monomer | 5FYL – antibody bound monomer complex | Incomplete glycans in PDB (NAG, BMA, AMA) | 26 | 30 | 1.4 |
| 10 | HIV-1 Env trimer | 5FYL – antibody bound trimer complex | Incomplete glycans in PDB (NAG, BMA, AMA) | 10 | 30 | 1.4 |
| 11 | HIV-1 Env trimer | One of PDBs from a MD trajectory | *N*-linked mannose-5 (BGA, BGL, BMA, AMA) | 23 | 30 | 1.4 |
| 12 | SARS-CoV-2 | One of PDBs from a MD trajectory | *N*-linked mannose-5 (BGL, BMA, AMA) | 23 | 30 | 1.4 |
| 13 | SARS-CoV-2 | One of PDBs from a different MD trajectory | *N*-linked mannose-5 (BGA, BGL, BMA, AMA) | 23 | 30 | 1.4 |
| 14 | SIVmac239 | One of PDBs from a MD trajectory | *N*-linked mannose-5, O-linked (BGA, BGL, BMA, AMA) | 23 | 30 | 1.4 |
| 15 | influenza HA | One of PDBs from a MD trajectory | *N*-linked mannose-5 (BGL, BMA, AMA) | 23 | 30 | 1.4 |
| 16 | influenza HA | H3N2 2020 homology model | *N*-linked mannose-5 (ASM) | 23 | 30 | 1.4 |
| 17 | Zika | One of PDBs from a MD trajectory | *N*-linked mannose-5 (BGLN, BMAN, AMAN) | 23 | 30 | 1.4 |
| 18 | Zika | One of PDBs from a different MD trajectory | *N*-linked mannose-5 (BGLN, BMAN, AMAN) | 23 | 30 | 1.4 |
| 19 | Adhesion domain of human CD2 | 1GYA | *N*-linked mannose-7 (NAG, BMA, MAN) | 23 | 30 | 1.4 |
| 20 | VSG3 | 6ELC | *O*-linked glycan (NAG, GLC) | 10 | 30 | 1.4 |
